# A novel generic dictionary-based denoising method for improving noisy and densely packed nuclei segmentation in 3D time-lapse fluorescence microscopy images

**DOI:** 10.1101/371641

**Authors:** Lamees Nasser, Thomas Boudier

## Abstract

Time-lapse fluorescence microscopy is an essential technique for quantifying various characteristics of cellular processes,i.e. cell survival, migration, and differentiation. To perform high-throughput quantification of cellular processes, nuclei segmentation and tracking should be performed in an automated manner. Nevertheless, nuclei segmentation and tracking are challenging tasks due to embedded noise, intensity inhomogeneity, shape variation as well as a weak boundary of nuclei. Although several nuclei segmentation approaches have been reported in the literature, dealing with embedded noise remains the most challenging part of any segmentation algorithm. We propose a novel denoising algorithms, based on sparse coding, that can both enhance very faint and noisy nuclei but simultaneously detect nuclei position accurately. Furthermore our method is based on a limited number of parameters,with only one being critical, which is the approximate size of the objects of interest. We also show that our denoising method coupled with classical segmentation method works properly in the context of the most challenging cases. To evaluate the performance of the proposed method, we tested our method on two datasets from the cell tracking challenge. Across all datasets, the proposed method achieved satisfactory results with 96.96% recall for *C.elegans* dataset. Besides, in *Drosophila* dataset, our method achieved very high recall (99.3%).

## Introduction

In cellular and molecular biology research, automatic segmentation and tracking of biological structures, i.e. cells or their nuclei are important tasks for further understanding of cellular processes. Time-lapse fluorescence microscopy (TLFM) is one of the most appreciated imaging techniques which can be used to quantify various characteristics of cellular processes, such as cell survival^1^, proliferation^2^, migration^3^, and differentiation^4^. The quantification of these processes plays a significant role in studying embryogenesis, cancer cells, stem cells, and other applications in the fields of molecular and developmental biology.

In TLFM imaging, not only spatial information is acquired, but also temporal information as well as spectral information, that produces up to *five-dimensional* (X, Y, Z + Time + Channel) images. Typically, the generated datasets consist of several (hundreds or thousands) images, each containing hundreds to thousands of objects to be analysed^5^. These large volumes of data cannot easily be parsed and processed, via visual inspection or manual processing within any reasonable time.

Nowadays, there is a growing consensus that automated cell segmentation methods are necessary to manage the time issue and provide a level of reliability and validity. Accordingly, the implementation of automated high-throughput cell nuclei detection and segmentation techniques may be able to improve the clinical diagnosis, predict the treatment outcome, and help to enhance therapy planning.

Although TLFM imaging is a very powerful technique and can capture valuable information from cellular structures, it still poses several limitations that can be summarised as follows: 1) non-uniform background illumination because of the fluorescence in cytoplasm and mounting medium; 2) low contrast and weak boundaries of non-obvious nuclei; 3) the degradation of image intensity over time due to photo-bleaching of fluorophores^5^. As a result of these limitations, obtained images become very noisy and difficult to interpret, which might lead to false detection and segmentation results.

Over the last few years, several methods have been proposed for filtering and denoising of cell nuclei microscopy images. Darbon *et al*.^6^ have used a Non-Local Mean approach to reduce the noise in microscopy images. This method relied on replacing the intensity value of a pixel with the average intensity values of the most similar pixels in the image. The algorithm demonstrated good performance in enhancing particles contrast, and reducing the noise in electron cryo-microscopy images. For Transmission Electron Microscopy (TEM) images, several digital filters have been introduced by Kushwaha *et al*.^7^ such as median or Wiener filter. A similar work proposed by Sim *et al*.^8^ and Aguirre^9^ based on employing an adaptive Wiener filter to enhance the effectiveness of the classical Wiener filter by considering the noise variance. Luisier *et al*.^10^ have suggested a *Poisson Unbiased Risk Estimation-Linear Expansion of Thresholds (PURE-LET)* technique for denoising images corrupted with Poisson noise. The method is based on three criteria: 1) minimising of an unbiased estimate of Mean Square Error (MSE) for Poisson noise, 2) linear parametrisation of the denoising process 3) preserving of Poisson statistics across scales. This algorithm is particularly promising for large datasets as well as images having a low signal to noise ratio. In addition, it has limited system requirements.

Over the past few years, deep learning methods have been successfully introduced for biological data processing. For instance, Liu *et al*.^11^ suggested a convolutional encoder-decoder neural network method for denoising of cell nuclei in fluorescence microscopy images. This method based on using the stochastic characteristics of noise as well as the shape of nuclei for learning step, and then regenerating the clean nuclei image based on the learned priori knowledge. Weigert *et al*.^12^ proposed a method to enhance the axial resolution in 3D microscopy images by reconstructing isotropic 3D data from non-isotropic acquisitions using a convolutional neural network. Another approach has also been proposed by Weigert *et al*.^13^ that presented a content-aware image restoration (CARE) networks method to denoise fluorescence microscopy data. This method introduced a solution to the problem of missing training data for deep learning in fluorescence microscopy by generating training data without the need for laborious manual annotations.

For cell nuclei segmentation literature, the reported approaches are classified into two categories: simple approaches such as thresholding method^14–17^, edge detection^18^ and shape matching^19–21^, and more sophisticated approaches like region growing^22–25^, energy minimization^26^ and machine learning^11,27–29^.

In the first category, R. Bise *et al*.^14^, Arteta *et al*.^15^, Liao *et al*.^16^ and Gul-Mohammed *et al*.^17^ applied a thresholding-based approach to segment cell nuclei. This approach assumes that the cell is usually brighter than its surrounding areas and there often exists an optimal threshold where individual cells can be segmented as separate objects. This assumption is not applicable in the challenging regions, because it is impossible to find a suitable threshold to separate all touching cells. Wählby *et al*.^18^ suggested to use an edge detection approach, in which an edge filter is applied to the image and therefore pixels are classified as edge or non-edge. These edges are usually detected by the first or the second order derivative method. Nevertheless, this method fails to detect the non-obvious cell’s boundary. Cicconet *et al*.^19^ and Türetken et al.^20^ proposed to use a shape matching-based approach. This approach depends on the assumption that cells, particularly nuclei, have round shapes. Hence, multi-scale blob detection^30^ can be employed to detect and segment cells.

In the second category, Cliffe *et al*^22^, Liu *et al*.^23^, Tonti *et al*.^24^ and Gul-Mohammed and Boudier^25^ employed region-based segmentation techniques in which the basic idea of these approaches is to combine the neighbouring pixels of initial seed points which have similar properties to form individual cells.

On the other hand, machine learning-based approaches can be implemented for cell segmentation. To give some examples, Ronneberger *et al*.^27^ proposed U-Net convolutional networks to segment cells by assigning class labels to every pixel in the image. A two-stage convolutional neural network method presented by Akram *et al*^28^ to precisely segment the cells. On the first stage, CNN is used to predict a regression on the cell bounding box. On the second stage, another CNN is employed to segment cells within the regressed bounding box. Liu *et al*.^11^ reported the use of a convolutional encoder-decoder network to segment the cell nuclei by learning stochastic characteristics of noise and shape of nuclei. Thus, segmented cell nuclei images can be generated. Moreover, Sadanandan *et al*^29^ advised to use the Deep Convolutional Neural Networks (DCNNs) to segment cells in fluorescence microscopy images. The aforementioned filtering and segmentation methods have achieved good results in microscopy images. However, most of these methods require tuning a set of parameters. Moreover, they may be used effectively only for specific applications. As a result, identifying a proper and robust approach for various datasets regarding high variation in cell nuclei volume, shape, and stain distribution as well as high cell density has become a major challenge in image analysis.

The contribution of this paper is to design a generic method for denoising of 3D cell nuclei images based on a sparse representation model. Consequently, a classical segmentation method can be used to segment cell nuclei without the need for more complicated segmentation approaches. The concept of dictionary learning, and sparse representation is already well established by M. Elad and M. Aharon^31^ as well as it is implemented in different application such as image denoising^32–34^, and image classification^35,36^. However, the added value of the proposed approach is employing the advantage of the sparse representation to find the potentials locations of nuclei in microscopy image. The novelty here is to obtain a denoised image and a detection map simultaneously. We believe that no similar studies have been reported in existing literature for denoising and simultaneously predicting objects location in images.

## Results

### Datasets description

Since the main motivation of our work is to automate the detection and segmentation of cell nuclei in time-lapse fluorescence microscopy images, we focused on applying our algorithm to development biology datasets, where only a limited number of existing methods had provided satisfactory results. The proposed framework is extensively tested on three real datasets for embryonic cells and one dataset of synthetic images with different values for the signal to noise ratio (SNR) and the object size. SNR is a performance measure for the sensitivity of imaging systems which defined as the ratio of the average signal level (*μ_signal_*) to the standard deviation (σ_*noise*_) of the background noise level: *SNR* = *μ_signal_*/σ_*noise*_. Expressed in logarithmic function as *SNR*(*dB*) = 10 × log (*μ_signal_*/¤_*noise*_).^37^

#### Synthetic dataset

In order to measure the robustness of the proposed method, we generated synthetic images of size (XYZ) equal (100 × 100 × 20) voxels containing spheres of two radii: 7 and 9 voxels. As it is common in fluorescence microscopy images to have low contrast and low signal to noise ratio (SNR) as a consequence of weak fluorescent staining or microscope properties. The images are distorted with different levels of Poisson-Gaussian noise, resulting in SNRs of 2 dB, −1 dB, −5 dB and −7 dB, respectively. Furthermore, the images include touching spheres where these conditions simulate the same characteristics exist in the real datasets as shown in Supplementary Fig 1.

#### Real dataset

The first dataset comes from the work of Gul-Mohammed^17^ and it is named as *CE-UPMC*. The other two datasets come from the cell tracking challenge^38,39^, and namely *Fluo-N3DH-CE* and *Fluo-N3DL-DRO*. The last two datasets are proven to be the hardest to be fully segmented automatically^38^. Each dataset from cell tracking challenge contains 2 sequences. For cell tracking challenge datasets, all pixels belonging to objects including the centroid are labelled as object by the ground truth. However, for the other dataset (i.e. CE-UPMC), only the centroid of each object is labelled. The datasets are described as follows:

***CE-UPMC dataset***. It involves *C. elegans* embryonic cells. The size (XYZT) of dataset is 512 × 512 × 31 × 160. The cells acquired every 1 minute using a spinning-disk confocal microscope. This dataset is very challenging, as the intensity of the images is decaying over time due to the labelling technique and acquisition system. Thus, the quality of the acquired images is low.
***Fluo-N3DH-CE dataset***. It includes *C. elegans* embryonic cells. The size (XYZT) of the first sequence is 708 × 512 × 35 × 250 and of the second sequence is 712 × 512 × 31 × 250. Both sequences are 8 – bit images with cells imaged every 1.5 minutes. The cells are acquired using a Zeiss LSM 510 Meta Confocal Microscope. This dataset is a challenging as well, since it has a low signal to noise ratio (SNR = 6.74 dB), in addition, the fluorescence can fade when the cells divide. Furthermore, the cells become smaller over time.
***Fluo-N3DL-DRO dataset***. It contains *Drosophila melanogaster* embryonic cells. The size (XYZT) of each sequence is 1272 × 603 × 125 × 49. Both sequences are 8 – bit images with cells imaged every 30 second. The cells are acquired using a SIMView light-sheet microscope. This dataset is very challenging as it has a large number of densely packed cells. In addition, it has a low signal to noise (SNR = 2.46 dB).

### Experimental setup and suitable parameters selection

Synthetic datasets are generated to study the effect of parameters (described in Table 1) on cell nuclei detection and segmentation, as well as to understand the overall mechanism for selecting and tuning the significant parameters of various datasets (summarised in Table 2).

**Table 1.** Description of denoising and segmentation parameters

| Parameters | Description |
| --- | --- |
| **Denoising** |  |
| Patch size [N N M] | The patch is a small region of an image with size ([N N M]). Patches are extracted by moving a window with a step size of one pixel over the raw image. |
| Dictionary size (K) | The dictionary is constructed by concatenating the patches to vectors (called atoms).<br>Dictionary size (K) is the total number of atoms. |
| Number of iterations (N) | A specified number of times to update the dictionary. |
| Sparsity level (L) | The number of nonzero elements (used atoms from the dictionary) for the sparse representation coefficient |
| **Segmentation** |  |
| SensitivityFactor | A scalar value within a range from zero to one.<br>It controls sensitivity towards thresholding more voxels as foreground. |
| NucleiSeedDilation | Radius of the structuring element for morphological dilation. |
| MinNucleiVolume | The approximate volume of the smallest cell nucleus in the image . |

**Table 2.** Denoising and segmentation parameters. When the values of parameters differ between the first and the advanced time points, the value for the advanced time points is given in round brackets.

| Parameters | Datasets |  |  |  |  |
| --- | --- | --- | --- | --- | --- |
|  | Synthetic | CE-UPMC | Fluo-N3DH-CE (seq1) | Fluo-N3DH-CE (seq2) | Fluo-N3DL-DRO(seq1 & seq2) |
| Patch size (voxels) | [15 15 5] | [ 20 20 5] | [25 25 5] | [25 25 5] | [10 10 5] |
| Threshold | global Otsu's<sup>44</sup> | local adaptive<sup>42</sup> | local adaptive<sup>42</sup> | local adaptive<sup>42</sup> | local adaptive<sup>42</sup> |
| SensitivityFactor | - | 0.56 (0.58) | 0.5 | 0.5 | 0.5 |
| MinNucleiVolume (voxels) | 20 | 5000 (1000) | 10,000 (3000) | 10,000 (3000) | - |
| NucleiSeedDilation (voxels) | 3 | 5 | 10 (5) | 20 (5) | 5 |

Our approach is based on the building of a dictionary (small patches of the image) that will be eventually used for denoising and detection of cell nuclei. We investigated the optimal size of the patches and the number of patches (called atoms) in the dictionary. Then, the randomly created dictionary is updated, and so we investigated the number of iteration for the update. Finally, we investigated the sparsity level (i.e., the number of used atoms) for the reconstruction of the denoised image and the detection map. To start with, we tested several values for patch size (p = 5 × 5 × 5, 10 × 10 × 5, 15 × 15 × 5 and 20 × 20 × 5), dictionary size (K = 64, 128, 256 and 512), sparsity level (L = 3, 6 and 9) and number of iterations (N = 5, 10, 15, 20, 25 and 30) at different noise levels (SNR = 2, −1, −5 and −7 dB). Since the coefficient of variation (CV) is a useful statistical descriptor for comparing the degree of variation from one data series to another one. The CV is defined as the ratio of the standard deviation to the mean^40^. Thus, we employed the CV to measure the effect of changing the parameters on the result. The average CV from the patch size, dictionary size, sparsity level and the number of iterations over the four noise levels are approximately 15, 2, 2 and 2 % respectively.

In all aforementioned parameters, we observed that the patch size is considered as a critical parameter where the change in this value has a major impact on the subsequent segmentation results as shown in Supplementary Fig. 2 (a). On the contrary, changes in the other parameters i.e. dictionary size, sparsity level and the number of iterations often achieve very close results as shown in Supplementary Fig. 2 (b), (c) and (d). Therefore, we fixed all parameters while tuned the patch size according to the object’s size present in the images.

In order to confirm the importance of patch size tuning, we conducted more analysis in the term of cell nuclei detection as shown in Supplementary Fig. 3. For example, at the first three noise levels 2, −1 and −5 dB all cell nuclei are correctly detected for different patch size values. However, at noise level equal −7 dB, many objects are falsely detected with the patch size equivalent top = 5 × 5 × 5 and one nucleus is not detected at patch size equal 20 × 20 × 5. Though, for patch size equal 10 × 10 × 5 and 15 × 15 × 5, all cell nuclei are correctly detected.

For all previously mentioned patch size values, the average of recall, precision, F-measure, and Jaccard index with different noise levels are presented in Supplementary Fig. 3 (b), where these measures are high when the patch size values equal p = 10 × 10 × 5 and 15 × 15 × 5 compared to the measures of the other two values.

Following the above experiment, we observed that, for robust detection and segmentation results, the patch size should not be less than 25% and not more than 100 % of the average cell nuclei volume in images.

### Denoising of 3D cell nuclei images

In this work, a sparse representation model^31,41^ is employed to obtain the denoised images. Our method is compared with the PURE-LET^10^, which is one of the most efficient, fast and automatic methods for denoising of multi-dimensional fluorescence microscopy images. The main motivation behind the need for cell nuclei denoising is assisting better segmentation of cell nuclei images. Therefore, the comparison between the denoising methods is performed in the context of improving segmentation results. For instance, the results in Supplementary Fig. 4 (first row), Supplementary Fig. 6 (a, c) and Supplementary Fig. 7 (a, c) show that our method is able to reduce, and almost remove the noise as well as enhance the contrast of cell nuclei. We have also noticed a better contrast than PURE-LET results as shown in Supplementary Fig. 4 (third row), Supplementary Fig. 6 (b, d) and Supplementary Fig. 7 (b, d).

For further assessment, thresholding-based approach is applied to the denoised images to obtain the segmentation mask. It can be noted from Supplementary Fig. 4 (second row), Supplementary Fig. 6 (e) and Supplementary Fig. 7 (e) that our method succeeded to segment all nuclei in comparison with the other method which failed to detect some nuclei as demonstrated in Supplementary Fig. 7 (f). Even though, PURE-LET method is able to detect all cell nuclei shown in Supplementary Fig. 6 (f), the size of segmented nuclei are smaller than their original size. Unfortunately, this method can not detect any cell nuclei at very low signal to noise ratios ( −5 dB and −7 dB,) as presented in Supplementary Fig. 4 (fourth row)

### Segmentation of 3D cell nuclei images

Following the denoising step, a local adaptive thresholding^42^ is applied to the denoised image to get the segmentation mask of candidates regions. In order to obtain the candidates locations of cell nuclei centres, we used a novel representation called the detection map. Each voxel in this map is computed as the summation of the patch coefficients that are used to reconstruct the denoised image. We then define a maximum response image by multiplying the denoised image with the detection map. This maximum response image is used to detect the local maxima (Fig. 1). Afterwards, the obtained local maxima are used as an input for a 3D marker-controlled watershed segmentation of the cell nuclei (Fig. 2).

**Figure 1.**
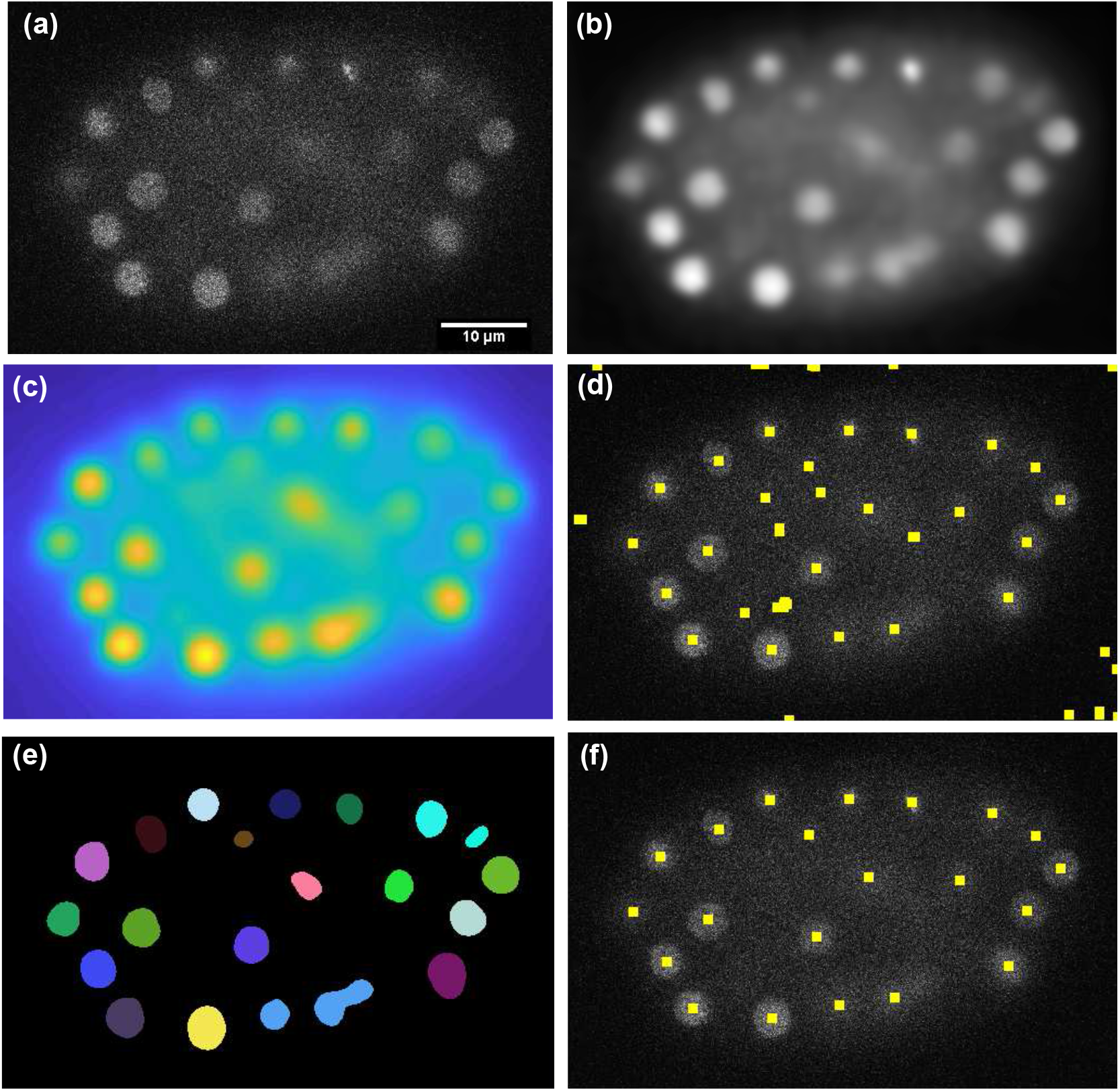
Denoising and nuclei detection with the sparse representation model. (a) A single plane (Z = 15) of time point (T = 100) from *CE-UPMC* dataset. (b) The denoised image obtained by applying the sparse representation model to the image in (a). (c) The detection map obtained from the sparse representation model for image in (a). (d) Marker points detected by applying the local maxima search on the maximum response image, obtained from multiplying image (b) with image (c). Marker points displayed as yellow squares are overlaid on the raw image. (e) Segmentation mask obtained by applying the initial segmentation to the image in (b). (f) Objects detected in the background are discarded by multiplying the detected marker points image (d) with the segmentation mask (e). Note that, the marker point detection here is performed in two dimensions for the purpose of explanation and visualisation, however, in the framework it is applied in three dimensions.

**Figure 2.**
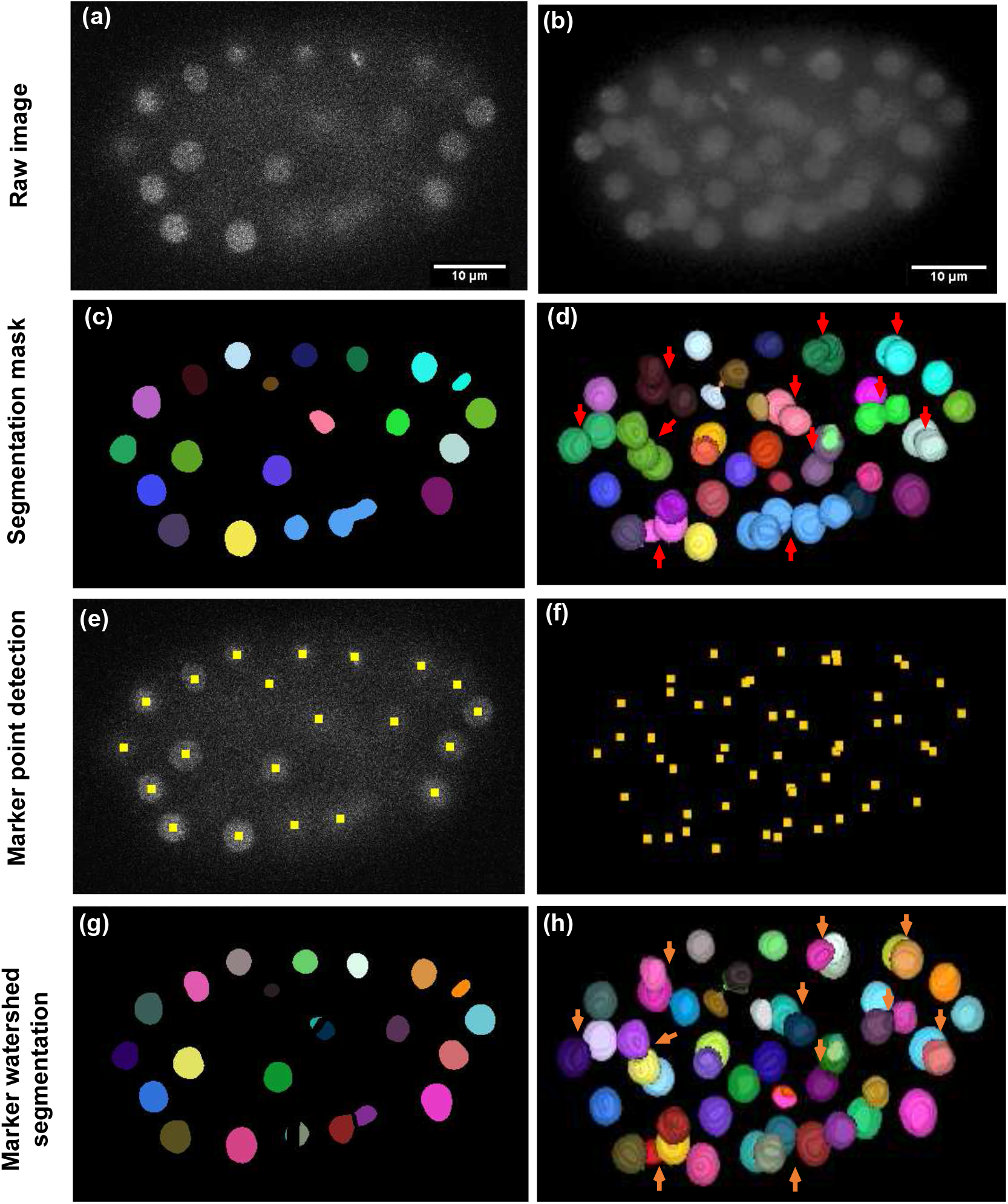
An overview of cell nuclei segmentation steps. First column: shows a single plane (Z = 15) of time point (T = 100) from *CE-UPMC* dataset. Second column: shows a three-dimensional view of the same time point. (a, b) The raw images. (c, d) The segmentation mask, which identifies the cell nuclei (presented as coloured) in the image, but fails to separate apparently touching cell nuclei (shown as red arrows). (e, f) Marker points (indicated by yellow squares) are obtained from the sparse representation model. (g, h) Marker-controller watershed segmentation that succeeds to separate apparently touching cell nuclei (orange arrows). Note that, different colours represent individual components. The marker points detection at (e) is performed in two dimensions for the illustration process. However, in the framework it is applied in three dimensions.

For the synthetic dataset, great performance is observed on very low signal to noise ratios (2 dB, −1 dB, −5 dB and −7 dB), in which our method is capable of correctly identifying and segmenting all cell nuclei at the various noise levels as presented in Supplementary Fig. 8. Furthermore, our method has similar performance as the top-ranked KTH algorithm from cell tracking challenge,^38,39^ as shown in Supplementary Fig. 5.

Regarding the *CE-UPMC* dataset, there is an intensity decay over time owing to the labelling technique and acquisition system. As a result, the acquired image quality is low. Table 3 and Supplementary Fig. 9, illustrate the results obtained at certain time points. For instance, at early time points (40, 60, 80 and 100) all cell nuclei are correctly detected. In addition, few false positives are also detected (2 objects out of 119 cell nuclei). Even though the image quality at advanced time points (120, 140 and 160) is low, only 11 cell nuclei out of 247 are not detected, also there exists a small number of false positives (3 objects out of 247 cell nuclei which is displayed as yellow arrows in Supplementary Fig. 9 (f). Typically, the reason for missing cell nuclei is the detection of clustered cell nuclei (indicated by red arrows in Supplementary Fig. 9 (f)) rather than detecting them separately. Supplementary Fig. 10 shows the segmentation results of our method and the results from the original paper^17^ of *CE-UPMC* dataset.

**Table 3.** Segmentation performance of our method (SRS) for *CE-UPMC* dataset.

| Time | GT | SN | TP | FN | FP | Recall (%) | Precision (%) | F-measure (%) |
| --- | --- | --- | --- | --- | --- | --- | --- | --- |
| **40** | 14 | 14 | 14 | 0 | 0 | 100 | 100 | 100 |
| **60** | 24 | 24 | 24 | 0 | 0 | 100 | 100 | 100 |
| **80** | 28 | 29 | 28 | 0 | 1 | 100 | 96.55 | 98.2447 |
| **100** | 51 | 52 | 51 | 0 | 1 | 100 | 98.08 | 99.03 |
| **120** | 57 | 53 | 53 | 4 | 0 | 92.98 | 100 | 96.3623 |
| **140** | 91 | 93 | 88 | 3 | 2 | 96.70 | 97.78 | 97.2370 |
| **160** | 104 | 101 | 100 | 4 | 1 | 96.15 | 99.01 | 97.5590 |

In the *Fluo-N3DH-CE* dataset, the proposed approach is able to identify and segment correctly more than 96% of total cell nuclei. Furthermore, it detects a small number of false positives (9 objects out of 876 cell nuclei) as well as a small number of false negative (29 cell nuclei). The achieved F-measure of approximately 97.8%, which is comparable to the competing algorithm, i.e. KTH^38,39^ (Table 4, Supplementary Fig. 11 and Supplementary Fig. 12).

**Table 4.** Segmentation performance of our method (SRS) and KTH^38,39^, for datasets from cell tracking challenge. The values shown in bold represent the highest performance. GT, number of cell nuclei in ground truth; SN, number of cell nuclei determined by the segmentation; TP, true positives; FN, false negatives; FP, false positives.

| Dataset | Algorithm | GT | SN | TP | FN | FP | Recall<sup>51</sup> (%) | Precision<sup>51</sup> (%) | F-measure<sup>51</sup> (%) | Jaccard<sup>52</sup> (%) |
| --- | --- | --- | --- | --- | --- | --- | --- | --- | --- | --- |
| **Fluo-N3DH-CE_01** | **SRS** | 478 | 464 | 459 | 19 | 5 | 96.03 | 98.9 | 97.44 | 66 |
| **Fluo-N3DH-CE_02** |  | 398 | 392 | 388 | 10 | 4 | 97.49 | 98.98 | 98.23 | 70 |
| **Total** |  | 876 | 856 | 847 | 29 | 9 | **96.69** | **98.94** | **97.84** | **68** |
| **Fluo-N3DH-CE_01** | **KTH** | 478 | 463 | 459 | 19 | 4 | 96.03 | 99.1 | 97.54 | 64 |
| **Fluo-N3DH-CE_02** |  | 398 | 394 | 386 | 12 | 8 | 96.98 | 97.97 | 97.47 | 59 |
| **Total** |  | 876 | 857 | 845 | 31 | 12 | 96.5 | 98.6 | 97.55 | 61.5 |
| **Fluo-N3DL-DRO_01** | **SRS** | 3792 | 122193 | 3757 | 35 | 118436 | 99.08 | 3.07 | 6 | 60 |
| **Fluo-N3DL-DRO_02** |  | 4097 | 120494 | 4082 | 15 | 116412 | 99.63 | 3.39 | 6.56 | 73 |
| **Total** |  | 7889 | 242687 | 7839 | 50 | 234848 | **99.37** | 3.32 | 6.43 | 66.5 |
| **Fluo-N3DL-DRO_01** | **KTH** | 3792 | 112535 | 3733 | 59 | 108802 | 98.44 | 3.32 | 6.4 | 62 |
| **Fluo-N3DL-DRO_02** |  | 4097 | 104655 | 4037 | 60 | 100618 | 98.54 | 3.86 | 7.4 | 78 |
| **Total** |  | 7889 | 217190 | 7770 | 119 | 209420 | 98.49 | **3.58** | **6.9** | **70** |

For *Fluo-N3DL-DRO* dataset, despite our method succeeds to detect more cell nuclei (99%recall), it has low precision (3%) due to the annotated ground truth, which considered only the cell nuclei located in the early nervous system and all other nuclei are deemed as false positives. As a result, the obtained F-measure is low with an approximate value of 6.4% Table 4. Furthermore, our method has a comparable segmentation accuracy with KTH competing approach^38,39^ (Table 4, Supplementary Fig. 13 and Supplementary Fig. 14).

The results achieved by our method for the two datasets obtained from cell tracking challenge are compared with the top-ranked KTH algorithm^38,39^. KTH algorithm is chosen for the reason that, it presented the best overall performance in the challenge. This algorithm is mainly based on adopting the band-pass filter to detect and segment cell nuclei.

Regarding the KTH algorithm, some detected objects are actually noise and some cell nuclei are not detected. This is because the algorithm detected clustered cell nuclei instead of detecting them separately. For example, at time point 28 from *Fluo-N3DH-CE* (seq1) and at time point 106 from *Fluo-N3DH-CE* (seq2), KTH algorithm failed to resolve the fusion of two nuclei (as presented by the red arrows in Supplementary Fig. 12 (b). In contrast, our method succeeds to identify and segment each nucleus individually as shown in Supplementary Fig. 12 (a) and (c). For *Fluo-N3DL-DRO* dataset, although our method succeeds to detect more cell nuclei than KTH approach (Table 4 and Supplementary Fig. 14), the evaluation method considered those cell nuclei as false positives, due to the annotation method which considered only the cell nuclei located in the early nervous system.

We have found that, the proposed method is less sensitive to some parameters such as dictionary size (K), sparsity level(L) and number of iterations (N). All these parameters are being fixed for different datasets and experiments with the subsequent values K = 64, L = 3 and N = 15. However, the proposed method is more sensitive to fundamental parameters, i.e., such as patch-size, and in a less critical manner to SensitivityFactor and NucleiSeedDilation. As these parameters are easy to understand, this makes them easier to tune-up if needed. We need to stress that all parameters, except patch size, are quite robust, as we only need to use three sets of parameters for all datasets. The set of empirically determined parameter values being applied to the datasets are listed in the Table 2.

Concerning the *Fluo-N3DH-CE* dataset, although the average cell nuclei size in sequence (2) is slightly greater than the average size in sequence (1), we have decided to use the same parameter (i.e., patch size) for denoising of both sequences. As a result, we tuned “NucleiSeedDilation” to avoid detection of multiple local maxima for the same object as explained in section “Marker points detection”.

We have also presented a Supplementary Table 1 to show the detection and segmentation results among various datasets considering the patch size percentage (related to average cell nuclei volume).

As a willing to test the genericity of the algorithm, we have conducted additional experiments on real datasets coming from various tissues such as thymus tissue (provided by J. Sheridan, Walter and Eliza Hall Institute of Medical Research (WEHI)), lymphoid tissue (provided by JR. Groom, WEHI), and islets of Langerhans tissue (come from the work of Tran *et al*.^43^), where robust cell segmentation is still challenging. Despite the noisy and crowded environment, the obtained results from our method are quite encouraging as presented in Supplementary Fig. 15.

## Methods

This section introduces a novel method for denoising and detection of cell nuclei in 3D TLFM images based on a sparse representation approach^31,41^. The use of sparse signal representation is becoming popular in several fields such as face recognition^45^, image denoising^32–34^ and inpainting^46^, and image classification^35,36^. Indeed, natural images represent very sparse data, especially in biology where numerous instances of the same structure, i.e. cell or nucleus, are present in the image. Moreover, a dictionary-based approach is usually linked to unsupervised learning since the data itself can be used to learn the basis vectors to build a sparse representation matrix.

The sparse representation method (shown in Fig. 3) is implemented as described by M. Elad and M. Aharon^31^, we have only changed the construction of the initial dictionary as depicted in the following steps. Firstly, the patches are extracted by moving a window with a step size of one pixel over the raw image. For each extracted patch, pixels’ intensities are summed up. Then, the average intensity over all patches is calculated. Secondly, an initial dictionary is constructed by selecting random patches from extracted patches, in particular,those having intensities greater than the obtained average intensity. By doing that, we are ensuring the presence of cell nuclei patches in the initial dictionary. Thirdly, a technique based on K-clustering with singular value decomposition (K-SVD)^41^ is implemented to update and obtain the final dictionary. Fourthly, the updated dictionary is used to reconstruct the denoised image as well as the detection map that will be used for detection of cell nuclei.

**Figure 3.**
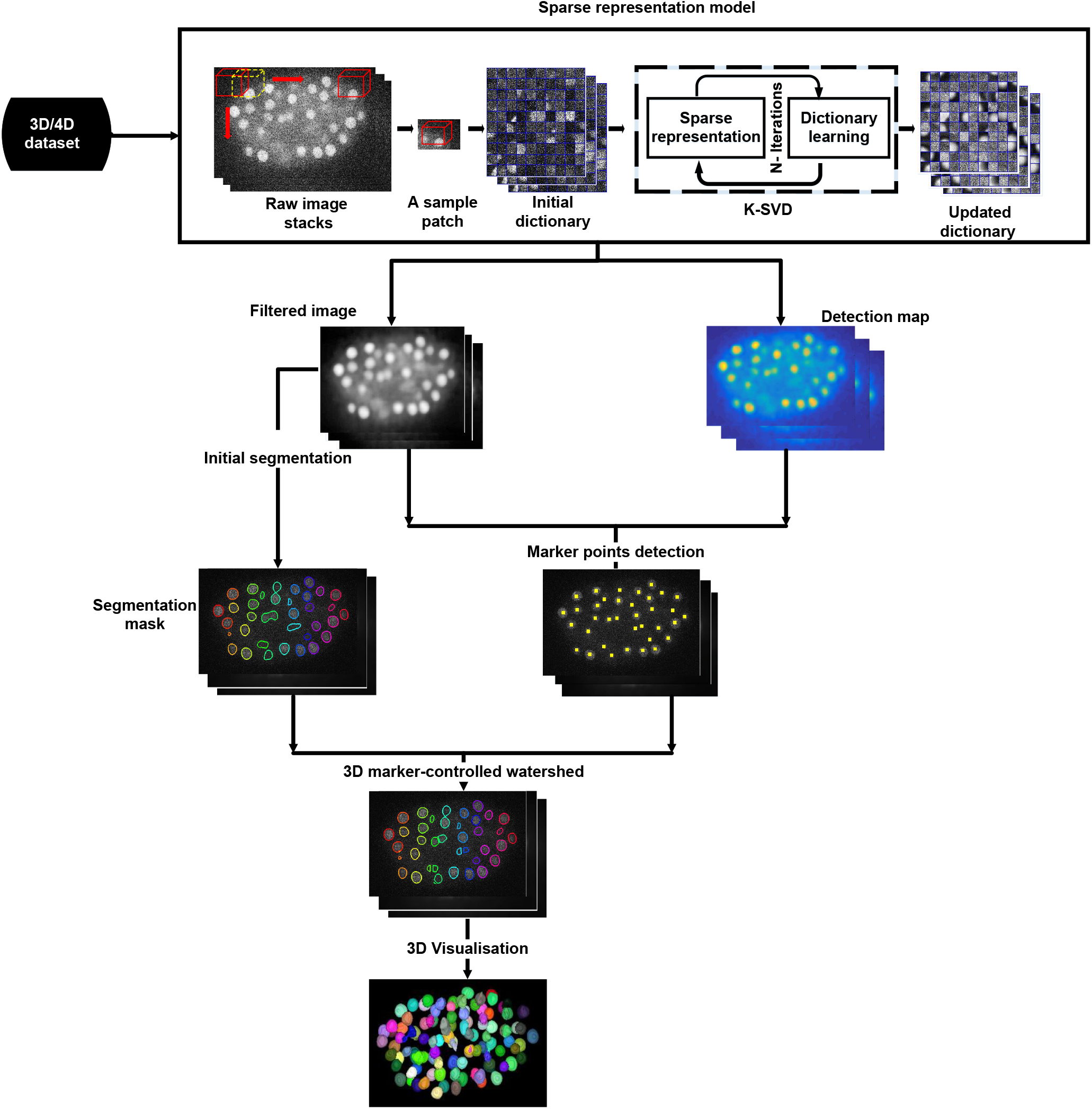
General representation of the proposed framework for denoising and segmentation of cell nuclei in 3D time-lapse fluorescence microscopy images. The proposed pipeline consists of data preprocessing, initial cell nuclei segmentation, cell nuclei detection, final segmentation as well as 3D visualization. In the preprocessing step, an initial dictionary is constructed by selecting random patches from the raw image as well as a K-SVD technique is implemented to update the dictionary and obtain the final one. Then, the maximum response image which is obtained by multiplying the denoised image with the detection map is used to detect marker points. Furthermore, a thresholding-based approach is proposed to get the segmentation mask. Finally, a marker-controlled watershed approach is used to get the final cell nuclei segmentation result and hence cell nuclei are displayed in a 3D view.

In the cell nuclei segmentation stage, the maximum response image, which is obtained by multiplying the denoised image with the detection map is used to detect the potential location of cell nuclei. Then, a thresholding-based approach^42,44^ is proposed to get the segmentation mask. Finally, a marker-controlled watershed approach^47^ is used to obtain the final cell nuclei segmentation result.

### An introduction to sparse representation

The idea of sparse representation is to obtain an efficient representation of a signal as a linear combination of few atoms chosen from a dictionary. Given a dictionary *D* ∈ *R^n×K^* that contains *K* atoms as column vectors *d_j_* ∈ *R^n^*, *j* = 1, 2,…, *K*. The sparse representation problem of a signal *x* ∈ *R^n^* can be described as finding the sparsest vector *α* ∈ *R^K^* where *X* ≃ *Dα*. The problem can be formulated as an energy optimization problem as follows:s

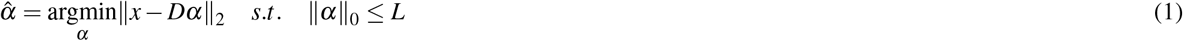

Where *x* is the signal, *α* denotes the sparse representation coefficients, ||*α*||_0_ is the *L*_0_ pseudo-norm that counts the number of non-zeros of *α* and *L* is a predetermined sparsity threshold.

Solving the previous optimisation problem is NP-hard and numerically intractable, thus several methods have been developed to get an approximate solution for this particular problem. The first type of methods uses L_0_ – norm minimisation, such as matching pursuit (MP)^48^ or orthogonal matching pursuit (OMP)^49^. The second type of method uses L_1_ – norm for optimisation. The objective of L_1_ – norm is to make the optimisation problem convex, which can be addressed efficiently using basis pursuit (BP)^50^.

The crucial issue in practical applications is to select the dictionary *D*. Basically, dictionaries are of two types: (1) fixed dictionaries, (2) adaptive dictionaries. The fixed dictionary as is the case for curvelet, discrete cosine, wavelet, ridgelet, or bandlet which use pre-defined and fixed atoms. This type of dictionary may not ensure a well-defined representation of all given signals. As a result, it is more appealing to use an adaptive dictionary approach to learn the dictionary directly from the data itself.

Learning the dictionary requires two-step, the first step is to compute an initial dictionary. It is usually computed by taking random patches directly from the raw image. These patches are overlapped with a step size of one pixel. To assure the presence of patches containing nuclei in the initial dictionary beside background patches, we select the patches whose intensity is greater than the average intensity of all patches extracted from the image. The second step is to update the initial dictionary by using the K-SVD algorithm^41^. This algorithm is a standard unsupervised adaptive dictionary learning algorithm that generalizes the well-known K-means clustering approach. It jointly learns a dictionary *D* = [*d*_1_, *d*_2_,…, *d_K_*] and a related sparse representation matrix *α* = [*α*_1_, *α*_2_,…, *α_m_*], *α* ∈ *R*^K^, from a set of training signals *X* = [*x*_1_, *x*_2_,…, *x_m_*], where each *x_i_* ∈ *R^n^* by solving the following problem:

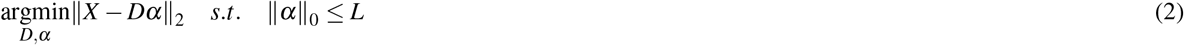

This technique solves the optimization problem by alternating between finding the sparse representation coefficients *α* and the dictionary D using an iterative approach. Assuming that *D* is known, the best sparse representation matrix is constructed by solving equation 2 using an orthogonal matching pursuit algorithm (OMP). Following the sparse representation stage, the representation vectors (*α*) are assumed to be fixed. Subsequently, the best dictionary is computed. Since finding the whole dictionary at the same time is impractical, the dictionary is updated atom by atom. Once the best dictionary and sparse representation coefficients are obtained, the denoised image and detection can be constructed.

#### Images with sparse representation

In this section, we present the reconstruction of the denoised image and detection map which will be used later in the detection and segmentation of cell nuclei

- **Denoised image reconstruction.** The denoised image with dictionary learning is formed by solving the following problem:

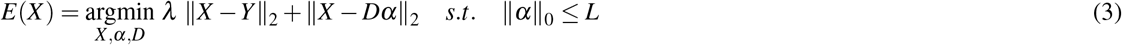 Where *X* is the ideal image, *Y* is the noisy image and *λ* is the regularisation parameter.
- **Detection map reconstruction.** In point of fact, the denoised image does not have sufficient contrast to completely separate touching nuclei. In order to improve cell nuclei detection, a detection map image that indicates the potential locations of cell nuclei will be built. The construction of this image is based on the computation of the sparse coefficients (*α_i_*) of each image patch. It can be obtained by:

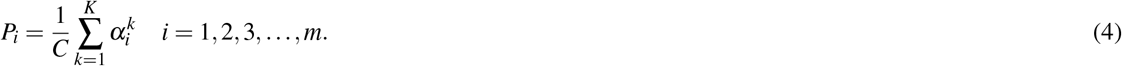 Where, *C* is a normalisation term, *P_i_* is the probability value corresponding to the *i* – *th* patch and 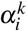 denotes the *k* – *th* element of *α_i_*. Notably, voxels within the centre of the nucleus have very high sparse coefficients values, in contrast to voxels far away from the centre having low values. Consequently, the *p_i_* value of the patches containing nuclei tend to be large compared with the *p_i_* value of background patches. As a result, dictionary learning technique with sparse representation can capture strong structures of biological images as well as restrain the noise.

### Cell nuclei segmentation

- **Initial cell nuclei segmentation.** A local adaptive thresholding approach^42^ is applied to the denoised image. The general concept of the algorithm is that for every image’s voxel the threshold is determined by the following equation:

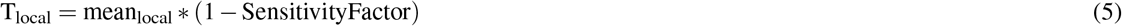

where, mean_local_ is the mean intensity value in the neighbourhood of each voxel and the SensitivityFactor is a scalar value within a range from zero to one which controls sensitivity towards thresholding more voxels as foreground. Accordingly, voxels with intensity values larger than T_local_ are set to 1, all others are set to 0. Small regions which detected as foreground and those smaller than a predefined volume denoted by (MinNucleiVolume) are discarded. This threshold corresponds to the volume of the smallest cell nucleus and is determined prior to the segmentation step. The resulting image is called the segmentation mask.
- **Marker points detection**. For splitting of touching cell nuclei, we employed a marker-controlled watershed technique. The marker points are obtained as follow: first, the denoised image is multiplied by the detection map to provide a maximum response image. Second, The maximum response image is processed to detect the local maxima (Fig. 4). The obtained local maxima image is multiplied by the segmentation mask to discard local maxima detected in the background. Third, a morphological dilation operator of certain radius denoted by (NucleiSeedDilation) is employed to avoid detection of multiple local maxima for the same object by merging those maxima that were in close proximity to each other. Finally, the modified image determining the marker points is fed to the subsequent watershed algorithm.
- **3D marker-controlled watershed segmentation**. Marker-controlled watershed segmentation is presented to separate connected cell nuclei clusters. The basic principle of watershed approach is to flood the denoised image, which contains merged objects starting at marker points as sources. Sometimes the flooding process is not stopped at the border of a cell nucleus, therefore the denoised image is multiplied by the segmentation mask prior to the flood. Eventually, watershed dams are built when different sources meet during the flooding process. This approach allows splitting clusters of apparently touching cell nuclei.

**Figure 4.**
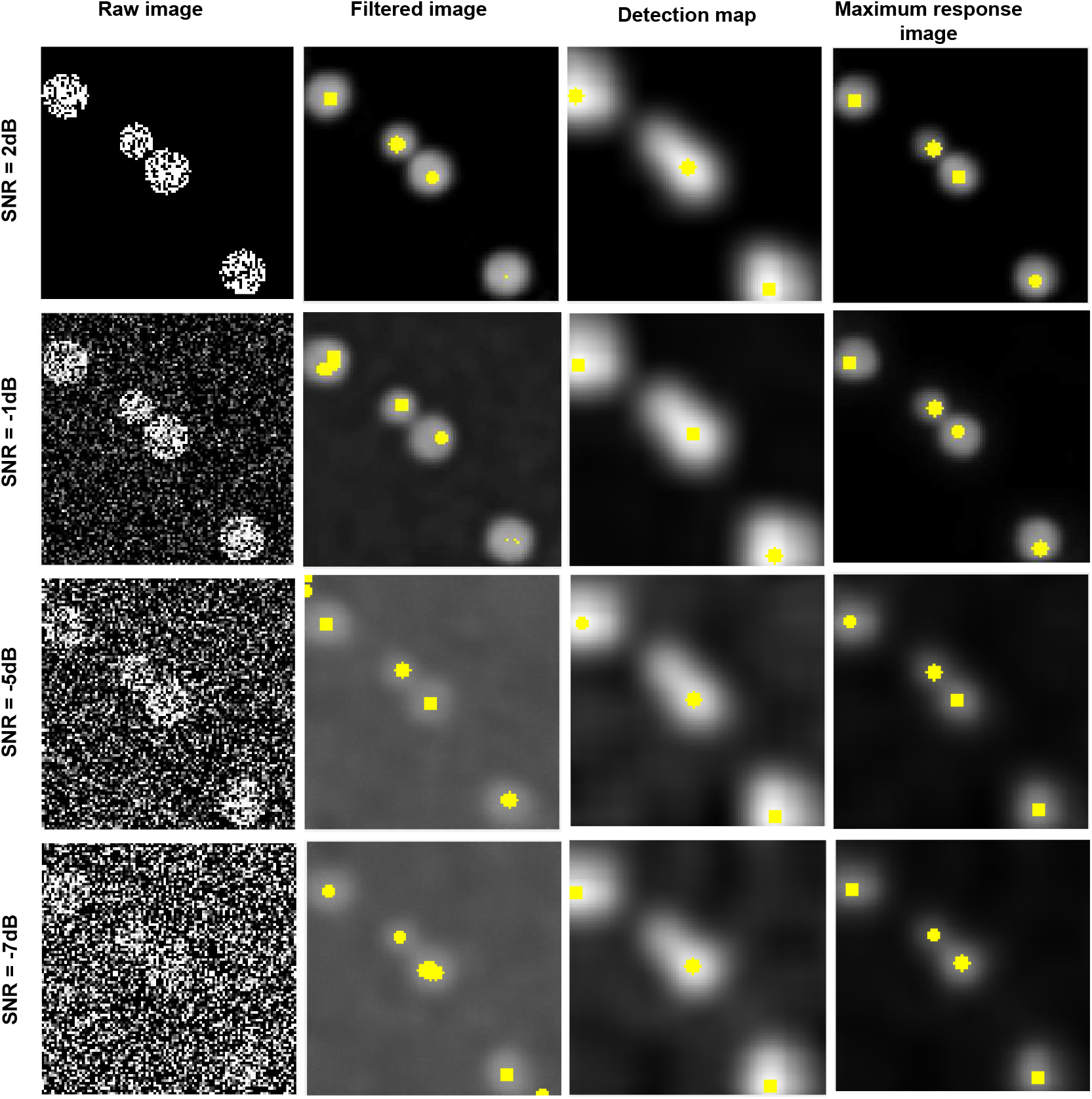
A comparison of marker points detection at various levels of noise. First column: representative single plane (Z = 10) of the raw image. Second column: the results of marker points detection from the denoised image. Third column: the result of marker points detection from the detection map. Fourth column: the result of marker points detection from the maximum response image. For all images the marker points depicted by yellow markers. Note that, the marker point detection here is performed in two dimensions for the purpose of explanation and visualisation. However, in the framework it is applied in three dimensions.

### Evaluation method and metrics

To assess the performance of the proposed algorithm, three metrics are employed. The first two metrics are the *recall*^51^ and *precision*^51^ of object detection. The recall is the proportion of the number of relevant detected cell nuclei to the total number of relevant cell nuclei in ground truth. Precision is the proportion of the number of relevant detected cell nuclei to the total number of irrelevant and relevant detected cell nuclei. These parameters are defined as follows:

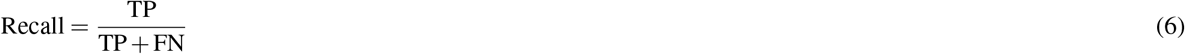

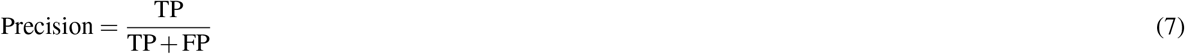

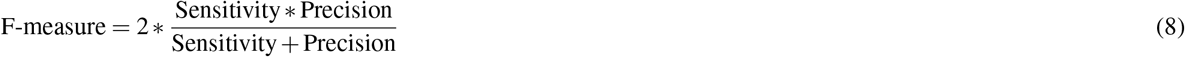

where, True Positive (TP) represents the total number of correctly detected nuclei, False Negative (FN) represents the number of undetected nuclei and False Positive (FP) represents the number of falsely detected nuclei. To compute these values, we used the following steps: first, we calculated the distance between the centroids of ground truth nuclei and centroids of segmented objects. Second, a weight is assigned to each pair of segmented and ground truth objects, equal to the distance between them. Third, Hungarian algorithm is used to solve this assignment problem. Objects with no match to any other object are considered as FP, objects absent in ground truth, but they appear in the segmentation result are deemed FP, and FN were objects absent in the segmentation result despite these objects appear in ground truth.

The third metric is the Jaccard index^52^ that measures the segmentation accuracy of the segmented objects. The Jaccard index for each set of segmented (A) and ground truth (B) objects is defined as the intersection between them divided by their union.

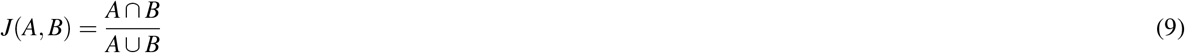

The final measure is then the average of the Jaccard indices of matched pairs.

### Implementation details

The image analysis framework is developed using MATLAB (R2017b) on a Windows-based computer (Intel Core i7, 3.07 GHz, and 16 GB RAM). Furthermore, the 3D ImageJ viewer plugin^54^ along with tools from the 3D ImageJ suite^53^ are used for three-dimensional visualisation of the final segmentation result. The source code, as well as datasets, are available upon request. Data processing using the complete framework took 5 mins for the synthetic dataset, 11 mins for the *CE-UPMC* dataset, 35 mins for *Fluo-N3DH-CE* dataset and 48 mins for *Fluo-N3DL-DRO* dataset to process only one time point of 3D image from the complete dataset. Regarding cell tracking challenge, the web site (http://www.codesolorzano.com/celltrackingchallenge) provides access to the datasets with the ground truth. In addition, it provides access to Windows and Linux executable files for the evaluation software as well as an executable program that includes the process description for KTH work.

## Conclusion

In this paper, we have presented a novel generic method for the denoising and detection of 3D cell nuclei in 3D time-lapse fluorescence microscopy images, based on a sparse representation model. We showed significant improvements over other denoising methods, and consequently, classical methods can be used for segmentation. We proposed, as to propose a complete workflow for denoising, detection and segmentation, to pair our denoising algorithm with a rather classical local thresholding methods and showed that we obtained similar or better results than state of the art algorithm. We observed than our denoising algorithm is performing extremely well for very noisy data and can hence help to detect very faint or previously undetectable nuclei. As the strength of our workflow is the denoising part, not so much the segmentation part, we observed (data not shown) little improvements for non-noisy data.

Over the last few years, deep learning approaches have achieved promising results in several domains, including denoising. However, they have some limits, for example, they are implemented to solve a specific problem, i.e., any new dataset will require a new training step. Furthermore, deep learning approaches required ground truth labels. Similarly to machine learning approaches, including deep learning, our method is based on a learning technique, but in dictionnary-based methods the learning is completely unsupervised and hence can be performed for any new data without any change.

We also showed that our algorithm can also lead to accurate detection of nuclei centroids, we coupled this detection to a classical segmentation method and showed very good results on challenging datasets. We believe than coupling our algorithm with a more powerful segmentation method may lead to even better results, but this was not the purpose of this article. We focused on a robust and powerful denoising method coupled with classical segmentation to provide a effective workflow with minimal tuning. The fundamental parameters which have a more noticeable impact on the result are patch-size, SensitivityFactor, and NucleiSeedDilation. All of these parameters are based on the average of cell nuclei volume present in the image. As these fundamental parameters are easy to understand, this makes them easier to tune-up if needed.

The obtained final segmentation results are quite good and stable. In addition, the training step is unsupervised and the dictionary can be directly learned from the image itself. We believe that no similar studies have been reported in existing literature for denoising and simultaneously predicting objects location in images. As a future work, we will investigate an online learning method to handle the time issue to reduce the processing time needed for dictionary learning.

The proposed method can handle the most challenging cases involving noisy, densely packed and multiple touching cell nuclei. In addition, it can produce the denoised image and simultaneously the potential locations of cell nuclei. The proposed method is adapted to the segmentation of cell nuclei in 3D time-lapse fluorescence microscopy images, nevertheless, it can be employed to detect and segment the nearly interacting intracellular organelles, including the endosomes, lysosomes, and lipid droplets. Our method is successfully evaluated on two embryo models, the *C. elegans*, and the *Drosophila* datasets. The overall detection and segmentation results are comparable to the existing methods, which is a good starting point for automated cell nuclei tracking process.

## Author contributions statement

L.N. wrote the manuscript text, analysed the data and developed the proposed algorithm; T.B. contributed to the conception and reviewed the manuscript.

